# A Label-Free Multi-Metric Pipeline for Benchmarking Single-Cell RNA-Sequencing Clustering and Testing the Reproducibility of Cell-Type Heterogeneity

**DOI:** 10.64898/2026.07.17.739160

**Authors:** Zachary Yousef, Jonah Simone, Dylan Klein, Heejoo (Jen) Cho, Avinav Bhuyan, Priya Bhatt, Jiawei Wu, Hongyi Wang, Li Cai

## Abstract

A discovered sub-population from single-cell transcriptomic data is only meaningful if it is reproducible, yet clustering is usually done with one method on one embedding and rarely tested. We present a label-free, multi-metric pipeline that reframes clustering as an auditable, methods-blind decision and separates two notions of stability that are commonly conflated: reproducibility under cell resampling (bootstrap) and reproducibility under re-embedding (retraining the representation). The pipeline evaluates seven clustering configurations across cluster counts using five non-redundant quality metrics. As a whole-dataset control on a mouse retinal atlas, it recovers an eight-cell-type annotation at 96.3% accuracy (adjusted Rand index, ARI = 0.91) without labels. We then validate the discovery mode on two cell types with opposite ground truth. On bipolar cells, which have well-established subtypes, the pipeline accepts the sub-structure: across-embedding reproducibility rises with cluster number to a high plateau (mean pairwise ARI ∼0.93 near the ∼15 known bipolar subtypes), with quality metrics improving in parallel. On rod photoreceptors, treated as homogeneous, it rejects over-clustering: the metric-selected partition passes a bootstrap-stability check but is not reproducible when the embedding is retrained (mean pairwise ARI = 0.69), and the metrics do not improve with cluster number. On synthetic data, the test recovers real structure down to a 5% subpopulation while rejecting null data (high sensitivity and specificity).

Bootstrap stability alone is therefore insufficient evidence for sub-population; the across-embedding test discriminates real sub-structure from over-clustering and applies to any cell type as a reproducible alternative to single-method, single-embedding clustering.

## Introduction

Single-cell RNA sequencing (scRNA-seq) has transformed the study of cellular diversity by profiling gene expression at the resolution of individual cells (Tang et al. 2009; Macosko et al. 2015). Whereas bulk RNA-seq averages expression across heterogeneous cells, scRNA-seq exposes the full spectrum of transcriptional states within a tissue and can reveal rare populations and within-cell-type heterogeneity that bulk methods cannot detect (Zheng et al. 2017; Svensson et al. 2018).

Identifying meaningful subtypes is a central analytical task, and many unsupervised clustering algorithms are available: centroid-based methods such as K-means (Lloyd 1982) and Gaussian mixture models (GMMs) (Scrucca et al. 2016), graph-based community detection such as Louvain and Leiden (Blondel et al. 2008; Traag et al. 2019), and deep representation learning such as the single-cell Decomposition using Hierarchical Autoencoder (scDHA) (Tran et al. 2021). Each method makes different geometric assumptions, and each can be run for any number of clusters k. The chosen method and k strongly influence biological conclusions, yet most analyses commit to a single configuration without systematic comparison (Duo et al. 2018; Freytag et al. 2018; Kiselev et al. 2019; Krzak et al. 2019).

How cluster quality is evaluated matters as much as how clusters are generated. The silhouette width (Rousseeuw 1987) captures geometric separation but performs poorly on continuous, anisotropic scRNA-seq embeddings (Kiselev et al. 2019). An entropy-based cluster-purity metric (ROGUE) (Liu et al. 2020) quantifies transcriptomic homogeneity, and cluster-wise bootstrap stability (Hennig 2007) tests whether a cluster is reproducible under resampling. No single metric is sufficient: each captures a distinct failure mode, and optimizing one in isolation can produce clusters that score well on one axis while failing to reflect biological structure.

Here we present a generalizable pipeline that reframes scRNA-seq clustering as a cell-subtyping decision-support problem. Rather than designating one algorithm as “best,” the pipeline asks whether subtype-level structure exists in a cell population and, if so, at what granularity and how reproducibly. Seven configurations, spanning centroid (K-means), probabilistic (GMM), and graph-based (Louvain, Leiden) algorithms applied to either a linear principal-component-analysis (PCA) embedding (Jolliffe and Cadima 2016) or a nonlinear scDHA embedding (Tran et al. 2021), are evaluated across a range of cluster numbers on five non-redundant quality metrics and compared by cross-method consensus. Critically, the pipeline distinguishes two forms of reproducibility that are commonly conflated: stability under cell resampling (bootstrap) and stability under re-embedding (retraining the representation).

Several scRNA-seq clustering benchmarks have been published, including pipeComp (Germain et al. 2020) and the systematic comparisons of Duo et al. (2018), Freytag et al. (2018), and Krzak et al. (2019). Our pipeline differs in three respects: it reports five non-redundant metrics separately rather than as a weighted composite, so the analyst can see what each metric eliminates and why; it is designed to operate label-free as a discovery tool for within-cell-type heterogeneity, where ground truth is unavailable by definition; and it integrates batch correction (Harmony) (Korsunsky et al. 2019) at both the whole-dataset and within-cell-type levels.

We demonstrate the pipeline in two settings (Fig. 1). As a positive control, we apply it to a whole retinal dataset with known cell-type structure and ask whether it recovers that structure without labels. As a discovery application, we apply it to the rod photoreceptors of the mouse retina (the most abundant retinal cell type, historically treated as homogeneous) and ask whether reproducible sub-states exist. Integrating six publicly available retinal scRNA-seq datasets spanning postnatal days P15–P91, we isolate the rod compartment and use the pipeline’s stability metrics to separate reproducible biology from over- clustering.

**Figure 1.**
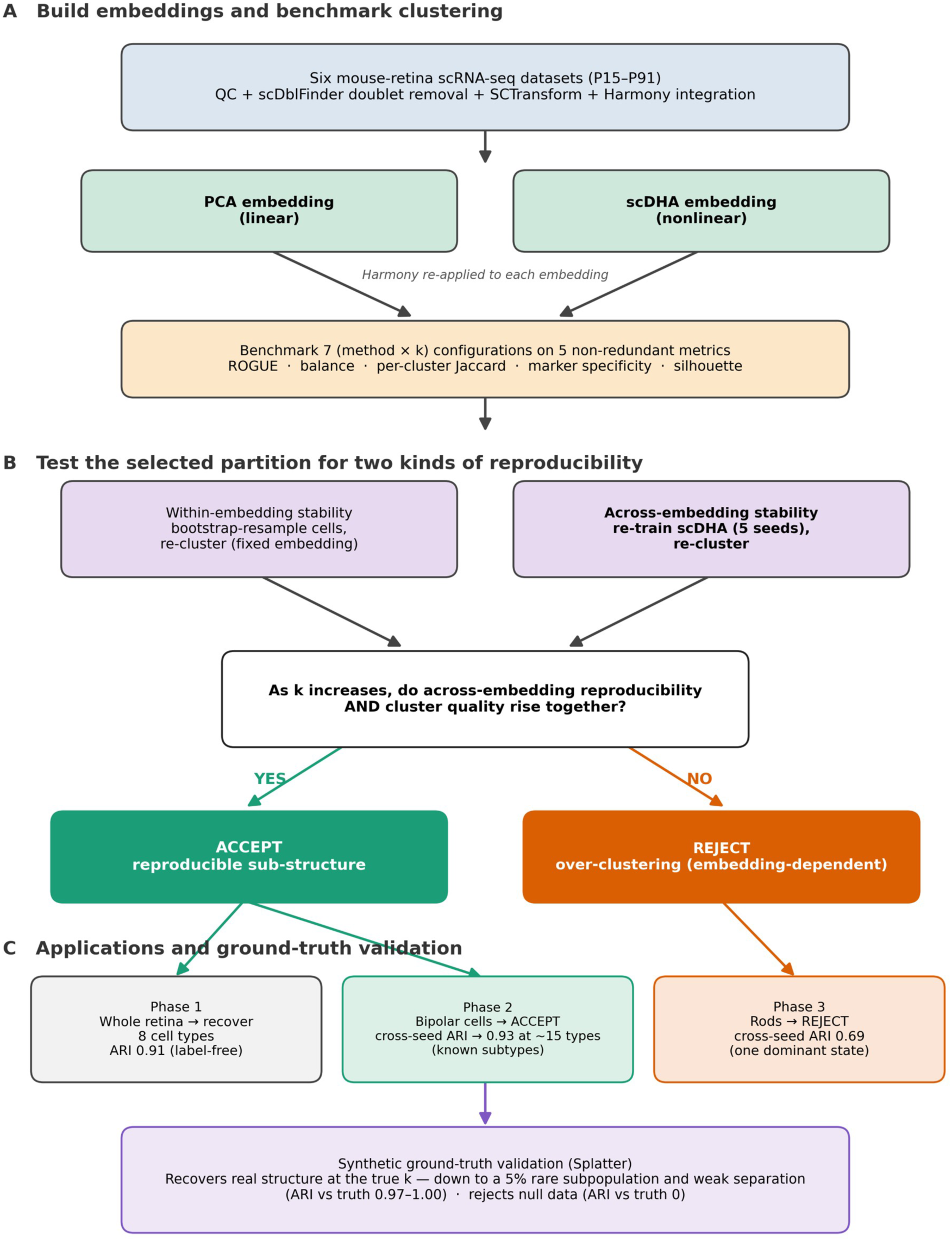
Pipeline overview and the reproducibility framework. (A) Six mouse-retina scRNA-seq datasets are QC-filtered, doublet-removed (scDblFinder), SCTransform-normalized, and Harmony-integrated; two embeddings, linear PCA and nonlinear scDHA, are built and Harmony-corrected, and seven (method, k) configurations are benchmarked on five non-redundant cluster-quality metrics (ROGUE, cluster-size balance, per-cluster Jaccard, marker specificity, silhouette). (B) The metric- selected partition is tested for two kinds of reproducibility: within-embedding stability (bootstrap resampling of cells in a fixed embedding) and across-embedding stability (re-training scDHA with five random seeds). The pipeline accepts a partition as reproducible sub-structure only if across-embedding reproducibility and cluster quality rise together as the cluster number increases; otherwise, the partition is rejected as embedding-dependent over-clustering. (C) The framework is applied to three retinal cases: whole-dataset cell-type recovery (Phase 1), bipolar cells (Phase 2, accepted), and rod photoreceptors (Phase 3, rejected); and validated against synthetic data with known ground truth.

## Results

### An integrated retinal atlas and label-based cell-type annotation

We assembled six publicly available mouse-retina scRNA-seq datasets spanning four developmental timepoints (P15, P49, P75, P91). After per-sample doublet removal with scDblFinder (Germain et al. 2021) and quality-control (QC) filtering, we retained 56,279 high-quality cells across 20,931 genes. Following normalization with SCTransform (Hafemeister and Satija 2019; Choudhary and Satija 2022) and integration with Harmony (Korsunsky et al. 2019), Louvain clustering at resolution 0.6 produced 34 fine-grained clusters, which canonical retinal markers collapsed into eight major cell types: rods (36,832 cells; 65.45%), bipolar cells (7,791; 13.84%), Müller glia (5,285; 9.39%), cones (2,873; 5.10%), retinal ganglion cells/neurons (2,279; 4.05%), retinal pigment epithelium (RPE; 520; 0.92%), amacrine cells (429; 0.76%), and microglia (270; 0.48%) (Fig. 2). The rod-dominant composition consists of prior mouse-retina atlases (Macosko et al. 2015; Shekhar et al. 2016). This label-based annotation provides the ground truth against which the label-free pipeline is scored below; the pipeline itself never uses it to select a partition.

**Figure 2.**
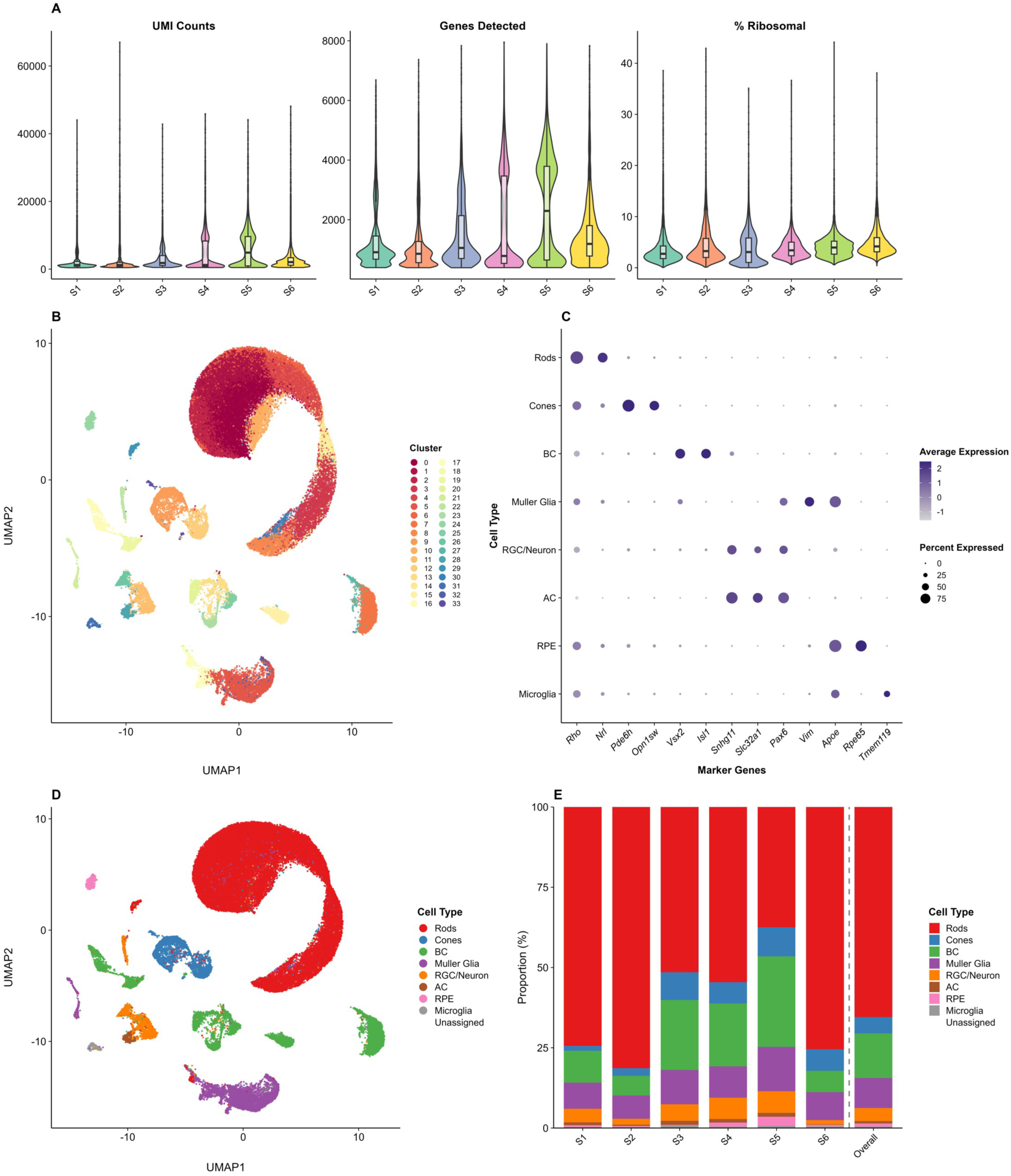
Quality control, integrated clustering, and cell-type annotation. (A) Violin plots of per-sample QC metrics on the analysis dataset: UMI counts, genes detected, and percent ribosomal. Cell-level filtering was applied upstream on the raw data using UMI-count, gene-count, percent-mitochondrial and percent-hemoglobin thresholds (Methods); mitochondrial and hemoglobin genes are removed after filtering, so percent ribosomal is shown here as the retained contamination metric. (B) Integrated uniform manifold approximation and projection (UMAP) of 56,279 cells colored by Louvain cluster (resolution 0.6). (C) Dot plot of canonical cell-type markers. (D) UMAP colored by the eight-type annotation. (E) Cell-type proportions across the six samples, with the aggregate composition at right. Rods are the majority in every sample.

### Phase 1 positive control: the pipeline recovers cell-type structure without labels

To test whether the pipeline recovers known biology, we ran all seven configurations on the Harmony-corrected whole dataset (n = 56,279) across k = 2–12, scoring each partition on the five methods-blind metrics, and computed the adjusted Rand index (ARI) (Hubert and Arabie 1985) and normalized mutual information (NMI) against the eight-type annotation only as a post-hoc check (Fig. 3). At the metrics- selected k = 8, PCA + Louvain recovered the annotation at ARI = 0.91 and NMI = 0.79, with 96.3% of cells assigned to the cluster matching their manual label. Rods, cones, and RPE mapped cleanly one cluster to one cell type; the remaining clusters merged transcriptionally similar types (Müller glia with microglia; amacrine cells with ganglion cells/neurons) or split the heterogeneous bipolar population, consistent with known bipolar diversity (Shekhar et al. 2016). Across the k sweep, PCA + Louvain peaked at ARI = 0.93 near k = 6 (where biologically related types merge) and remained above 0.90 at k = 8. Comparing all methods at k = 8, PCA + Louvain led (ARI = 0.91), followed by scDHA + Louvain (0.73) and scDHA + Leiden (0.71); the centroid methods were capped at ARI ≤ 0.62 because their near-equal-sized clusters cannot recover the three rare cell types (<1% each), and PCA + GMM degraded because its elliptical-cluster assumption fits the rod-dominated geometry poorly. A conventional analysis would over-cluster to 34 groups and collapse them by hand; the pipeline reproduces the eight-type annotation through an automated, auditable, methods-blind benchmark without seeing a label, marker panel, or analyst decision.

**Figure 3.**
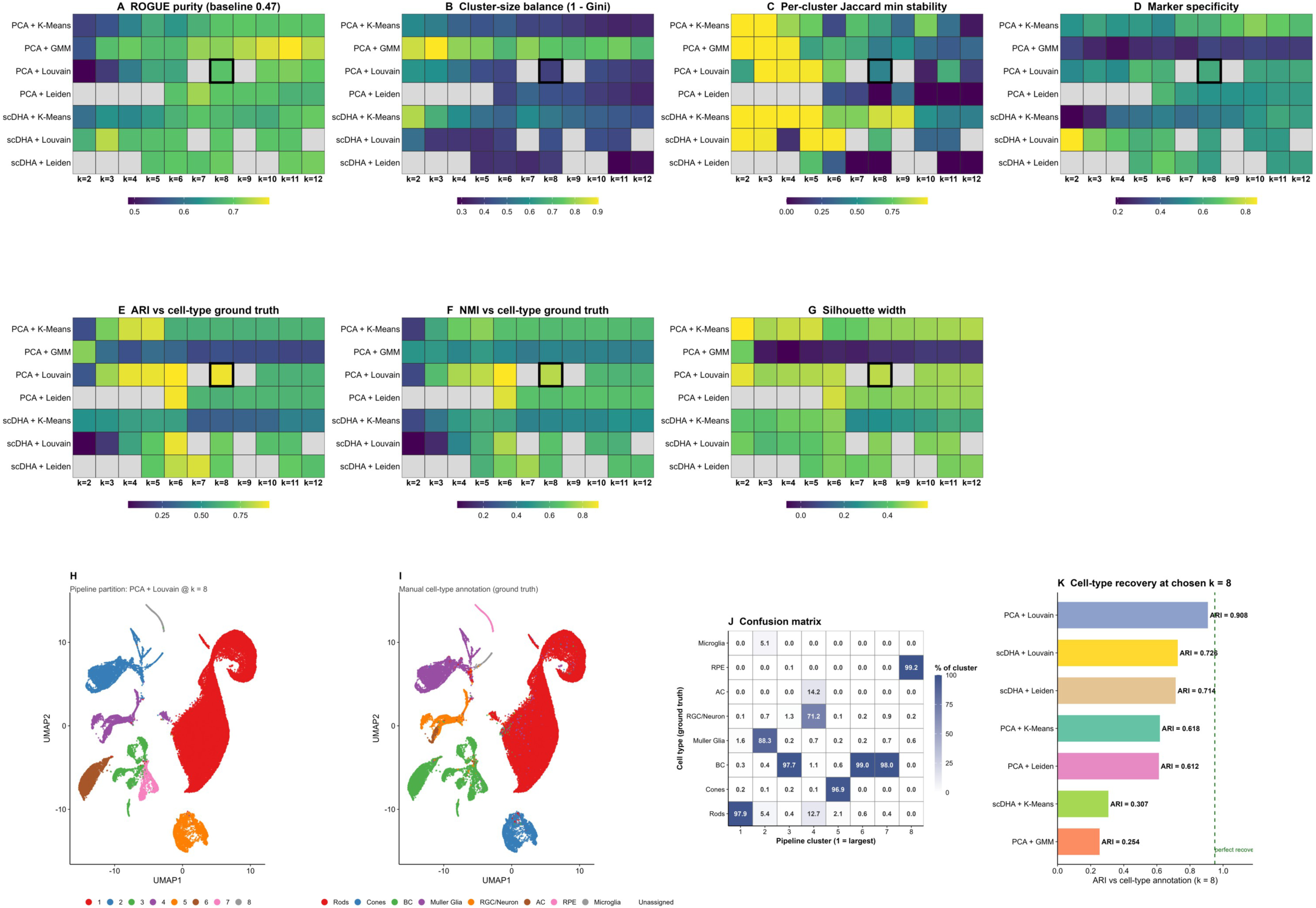
Phase 1 positive control on the whole dataset (n = 56,279). (A–G) Heatmaps of the seven configurations (rows) over k, showing the five methods-blind metrics plus the post-hoc ARI and NMI against the eight-type annotation. The chosen partition (PCA + Louvain, k = 8) is boxed; heatmap cells are colored by metric value, with exact per-configuration values tabulated in Supplementary Table S1. (H) UMAP of the pipeline partition; (I) the same coordinates colored by the manual annotation. (J) Cluster-by-cell-type confusion matrix at k = 8 (ARI = 0.91, NMI = 0.79, per-cell accuracy = 96.3%). (K) Cell-type recovery at k = 8 for every method, sorted by ARI; PCA + Louvain leads.

### Phase 2 positive discovery control: the pipeline accepts reproducible bipolar-cell subtypes

The Phase 1 control shows the pipeline recovers known cell types; a discovery tool must also correctly accept sub-structure that is genuinely present within a cell type. We therefore applied the discovery mode to the bipolar cell compartment (7,791 cells), which is known to comprise multiple reproducible transcriptomic subtypes (Shekhar et al. 2016). We trained five independent scDHA embeddings, swept the cluster number from k = 2 to 18, and measured across-embedding reproducibility (the mean pairwise ARI between the five resulting partitions) alongside the quality metrics (Fig. 4). In direct contrast to the rods below, bipolar sub-structure became progressively more reproducible as k increased toward the biological subtype count: the mean pairwise ARI rose from 0.19 at k = 2 to a high plateau of ∼0.93 (minimum pairwise ARI ∼0.90) across k ≈ 14–18, while silhouette width rose from 0.17 to ∼0.42 and ROGUE and cluster-size balance improved in parallel (Fig. 4B,C). Critically, this reproducibility plateau coincides with the ∼15 transcriptomic bipolar types reported for the mouse retina, so the across- embedding agreement is maximized precisely at the granularity where the known biology sits, not below it. This concordant rise of reproducibility and cluster quality toward the true count is exactly the signature the pipeline is built to detect. We take the k ≈ 15 partition, which the resolution search recovers as 14 clusters, as the accepted solution; it separates the bipolar compartment into 14 well-separated subclusters spanning 32.3% to 1.6% of the compartment, each forming a visually distinct island in the embedding (Fig. 4A,D), the direct visual counterpart to the single continuous rod manifold shown below. Bipolar cells thus provide positive discovery control (the pipeline accepts real sub-structure), complementing the whole-dataset annotation-recovery control of Phase 1.

**Figure 4.**
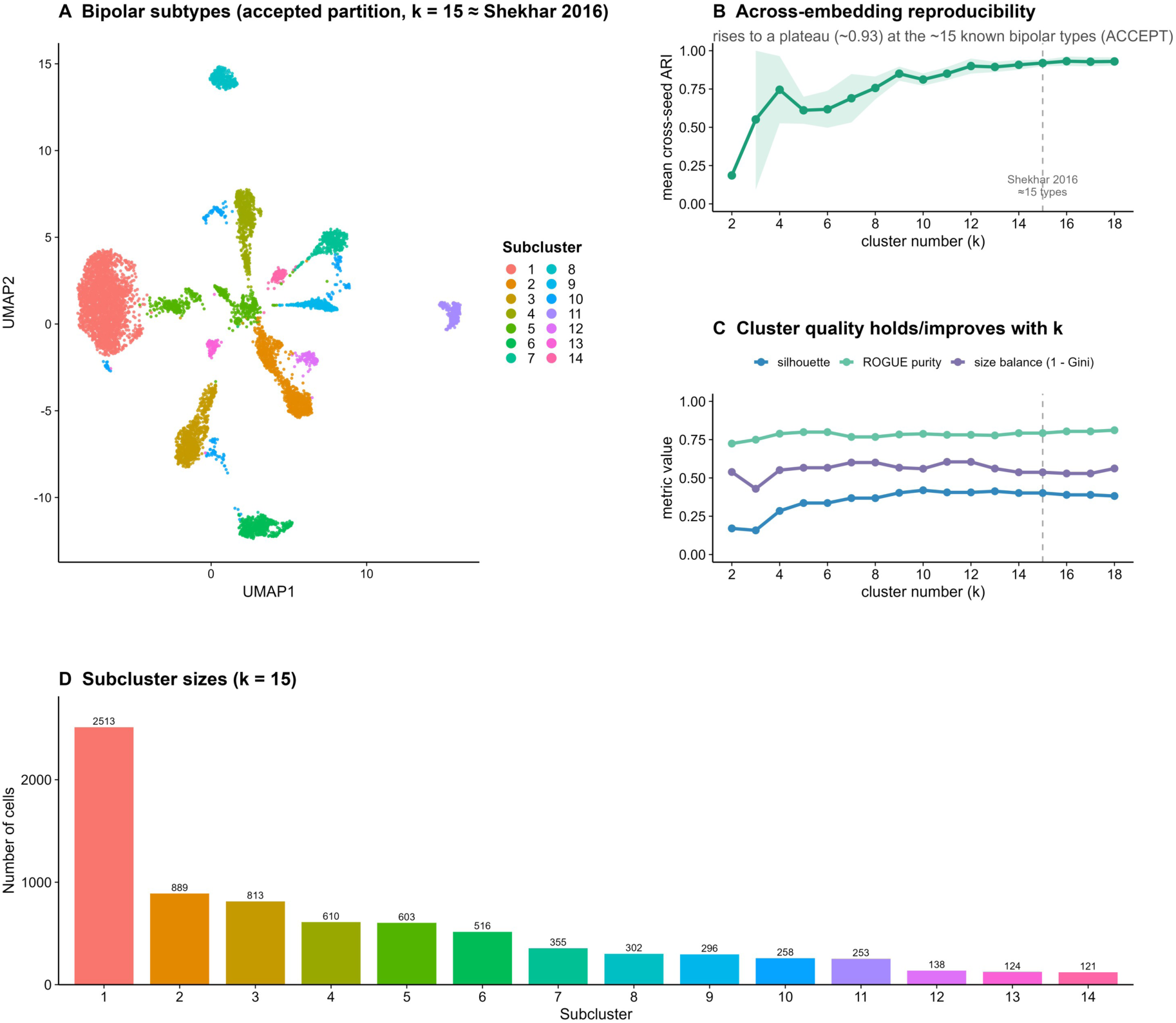
Phase 2 positive discovery control: the pipeline accepts reproducible bipolar-cell subtypes (n = 7,791). The discovery mode applied to the bipolar compartment, which has well-established transcriptomic subtypes, using five independently trained scDHA embeddings. (A) UMAP of the reference scDHA embedding at the accepted partition (target k = 15, recovered as 14 clusters, close to the ∼15 known bipolar types), colored by subcluster; the bipolar cells resolve into 14 well-separated islands, the visual opposite of the single continuous rod manifold (Figure 5). (B) Mean pairwise across- embedding ARI versus cluster number k (shaded band, min–max across the five embeddings); reproducibility rises to a high plateau, reaching ∼0.93 across k ≈ 14–18 (minimum ∼0.90) and peaking at the ∼15 known bipolar types, the signature the pipeline uses to ACCEPT sub-structure. (C) Cluster- quality metrics (silhouette, ROGUE purity, size balance) versus k, which hold or improve in parallel with reproducibility. (D) Subcluster sizes at k = 15, spanning 32.3% to 1.6% of the compartment. The concordant rise of reproducibility and quality (contrast the rods, Figure 5) is what distinguishes genuine sub-structure from over-clustering.

### Phase 3 negative case: the pipeline rejects non-reproducible rod over-clustering

We next applied the same seven configurations to the cone-cleaned rod subset (36,832 rods identified by *Rho* and *Nrl* expression; 36,771 retained after removing 61 cone-contaminated cells) across k = 2–9, computing parallel PCA and scDHA embeddings, re-applying Harmony to each, and scoring every partition on the five metrics (Fig. 5A–E). Each metric isolates a distinct failure mode. ROGUE confirmed that splitting the rods produced transcriptionally tighter groups than the unsplit population (baseline 0.79; all methods above it). Cluster-size balance (1 − Gini) separated the methods into two regimes: the Leiden configurations collapsed below 0.4 for k ≥ 3, whereas the centroid and Louvain methods produced usable partitions whose smallest cluster contained hundreds to thousands of cells. Per-cluster bootstrap Jaccard and marker specificity favored low k, and silhouette width, low across the field, as expected for continuous scRNA-seq embeddings (Kiselev et al. 2019), was highest for the graph methods at low k (scDHA + Louvain at k = 3: silhouette 0.33). Read together, the panel converges on a low-k solution, and the metrics select scDHA + Louvain at k = 3 as the representative partition, resolving the rods into a dominant cluster of 25,600 cells (69.6%), a second cluster of 7,607 cells (20.7%), and a smaller cluster of 3,564 cells (9.7%) (balance 0.60; ROGUE 0.89) (Fig. 5F).

**Figure 5.**
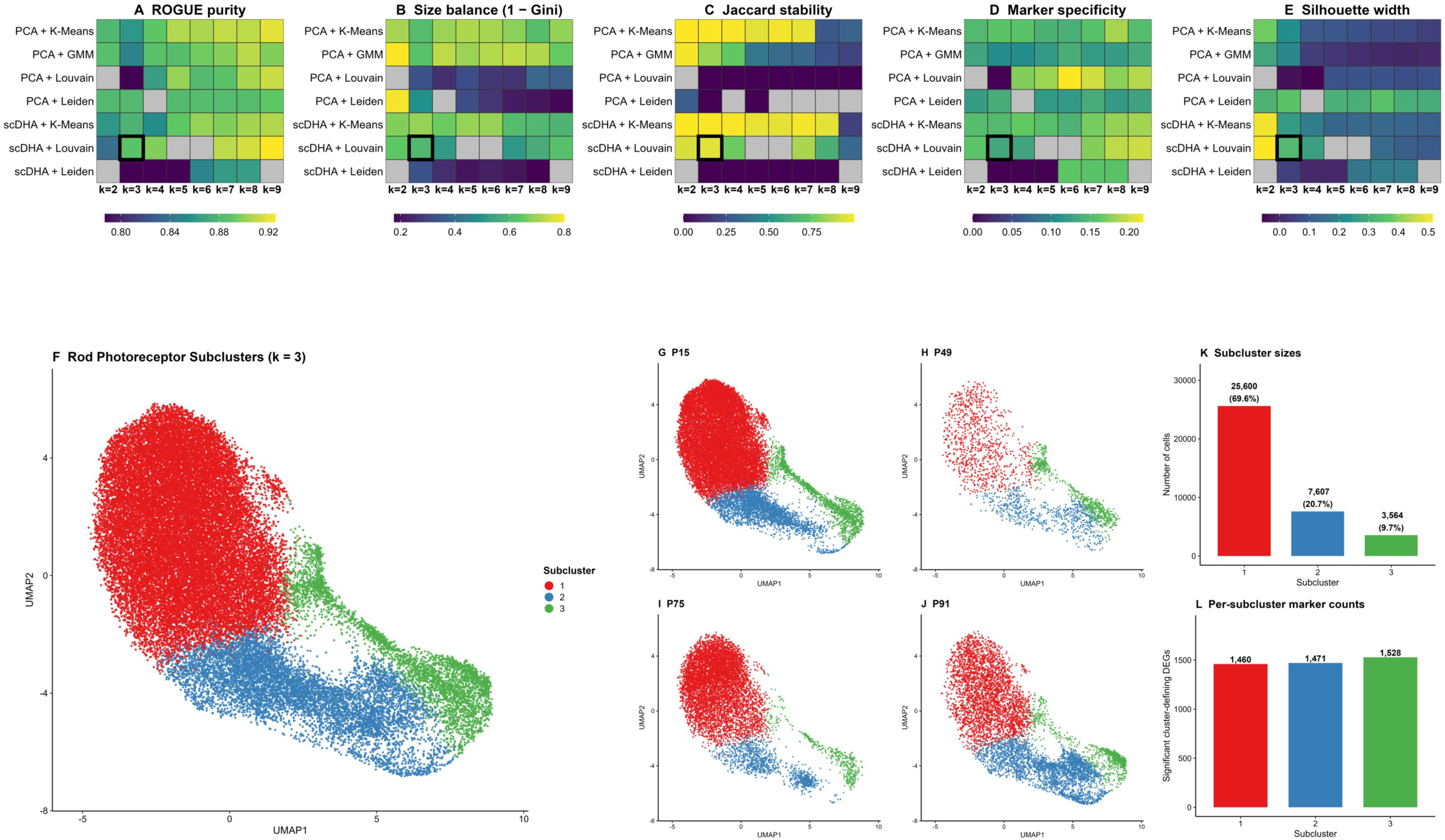
Phase 3 rod benchmark (n = 36,771) and the metric-selected partition. (A–E) Heatmaps over the seven configurations and k = 2–9 for the five metrics: (A) ROGUE purity; (B) cluster-size balance (1 − Gini); (C) minimum per-cluster bootstrap Jaccard; (D) marker specificity; (E) silhouette width. Higher is better for all five; the metric-selected scDHA + Louvain at k = 3 is boxed; heatmap cells are colored by metric value, with exact per-configuration values tabulated in Supplementary Table S2. (F) UMAP of the 36,771 rods colored by the k = 3 partition (sizes 25,600 / 7,607 / 3,564). (G–J) Per- timepoint UMAPs (P15, P49, P75, P91). (K) Subcluster sizes. (L) Number of cluster-defining DEGs per subcluster. Across-embedding reproducibility of this partition is evaluated in Figure 6 and Supplementary Figure S4.

### The dominant rod state is reproducible; the finer partition is not

We then tested whether the selected partition is a property of the rod data or of the particular embedding on which it was computed, using two complementary analyses (Supplementary Fig. S4). *Within a fixed embedding,* the k = 3 partition is highly stable to cell resampling: across 50 bootstraps (80% subsampling) the minimum per-cluster Jaccard was 0.95 and the mean partition-to-bootstrap ARI was 0.97. *Across embeddings,* however, the partition is not reproducible. Re-training scDHA with five independent random seeds on the identical input, varying only the stochastic embedding, yielded a mean pairwise ARI of 0.69 (range 0.51–0.81) between the resulting k = 3 partitions, and the smallest of the three clusters ranged from 134 to 5,149 cells depending solely on the training seed. In every case the dominant cluster (∼70% of rods) was preserved essentially intact, but the assignment of the remaining cells reshuffled from run to run. Because scDHA is stochastic even at a fixed seed, the specific minor-cluster identity recovered in any single run is not a stable feature of the data. The pipeline therefore reaches a clear, reproducible conclusion: mouse rod photoreceptors are dominated by a single transcriptional state that is robust across datasets, embeddings, and clustering parameters, whereas any finer partition reflects the chosen embedding rather than discrete biological sub-states. This result also demonstrates that within-embedding resampling stability, while necessary, is not sufficient evidence for a reproducible sub-population.

Interpreted positively, rod photoreceptors comprise a single dominant transcriptional state with continuous residual variation (the k = 2 split has the highest silhouette of any rod partition, 0.51, indicating real but graded variation) rather than a set of discrete, reproducible subpopulations.

### Across-embedding stability discriminates real from artifactual sub-structure

Placing the two discovery cases side by side isolates the pipeline’s core capability (Fig. 6). Both bipolar cells and rods pass a within-embedding bootstrap-stability check at their metric-selected partitions (minimum per-cluster Jaccard ≈ 0.9), so bootstrap stability alone cannot distinguish them. The across- embedding test does. As the cluster number increases, bipolar reproducibility climbs to a plateau at the true subtype count (mean pairwise ARI to ∼0.93 across k ≈ 14–18, coinciding with the ∼15 known bipolar types) with silhouette width rising in parallel (0.17 to 0.42), whereas rod reproducibility peaks at only ∼0.72 (its maximum, at k = 4; 0.69 at the metric-selected k = 3) and then declines while rod silhouette falls monotonically (0.51 to 0.11), forcing finer rod partitions degrades cluster quality without gaining reproducibility, the signature of over-clustering rather than of nested biological structure. The pipeline therefore accepts bipolar sub-structure and rejects rod over-clustering by a single automated criterion, whether across-embedding reproducibility rises together with cluster quality as granularity increases. This is the operational definition the pipeline provides for a real sub-population, and it is the property that neither single-embedding clustering nor bootstrap stability can supply on its own (Table 1).

**Figure 6.**
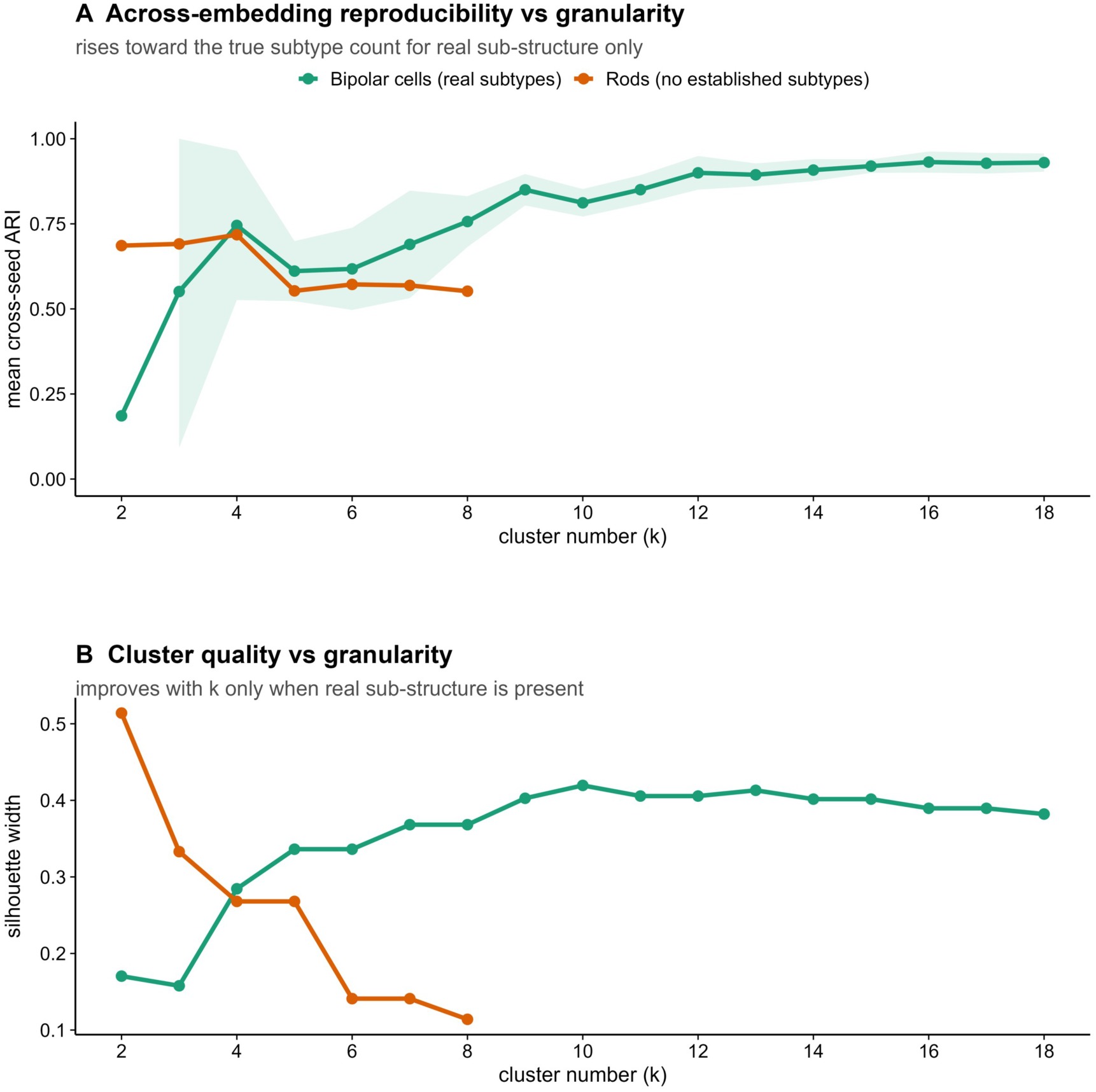
Across-embedding stability discriminates real from artifactual sub-structure. The discovery mode applied to two cell types with opposite ground truth, using five independent scDHA embeddings per cell type. (A) Mean pairwise across-embedding ARI versus cluster number k for bipolar cells (known subtypes) and rods (no established subtypes); the shaded band (bipolar) shows the min–max pairwise ARI across the five embeddings. Bipolar reproducibility rises to a plateau at the true subtype count (to ∼0.93 across k ≈ 14–18, coinciding with the ∼15 known types), whereas rod reproducibility peaks at only ∼0.72 (its maximum, at k = 4; 0.69 at the metric-selected k = 3) and then declines. (B) Silhouette width versus k for the two cell types, rising for bipolar cells and flat/declining for rods, the cluster-quality metrics move in concordance with reproducibility only when real sub-structure is present. Together these panels show the pipeline accepting bipolar sub-structure and rejecting rod over-clustering by the same automated criterion.

**Table 1.** Summary of the pipeline’s three demonstrations on the mouse retinal atlas. Across-embedding reproducibility is the mean pairwise adjusted Rand index (ARI) between partitions from five independent scDHA embeddings.

| Demonstration | Cell population<br>(n) | Ground truth | Metric-selected k | Across-embedding<br>reproducibility | Pipeline verdict |
| --- | --- | --- | --- | --- | --- |
| **Phase 1 —**<br>**whole-dataset**<br>**control** | Whole retina<br>(56,279) | 8 annotated cell<br>types | 8 | ARI 0.91 vs.<br>manual annotation | Recovers cell<br>types without<br>labels |
| **Phase 2 —**<br>**positive**<br>**discovery control** | Bipolar cells<br>(7,791) | ~15 established<br>subtypes | ~15 | cross-seed ARI<br>rises to ~0.93<br>(plateau at ~15<br>types) | Accepts real sub-structure |
| **Phase 3 —**<br>**negative case** | Rod<br>photoreceptors<br>(36,771) | No established<br>subtypes | 3 | cross-seed ARI<br>0.69, then<br>declines | Rejects over-clustering |

### Ground-truth validation on synthetic data

To confirm on ground truth what the bipolar/rod comparison showed on real data, we validated the pipeline on synthetic scRNA-seq datasets simulated with Splatter (Zappia et al. 2017) (Fig. 7). We generated null datasets containing a single homogeneous population and structured datasets containing three groups at varying separation (differential-expression probability de = 0.03–0.30), including a rare- subpopulation case in which one real group comprised only 5% of cells. For every structured dataset the pipeline recovered the true partition at the true cluster number: the ARI against ground truth reached 0.97 (weak, de = 0.03), 0.999 (medium), and 1.00 (strong, and the rare 5% subpopulation) at k = 3, and across- embedding reproducibility peaked at the same k (Fig. 7A,B). For the null datasets the pipeline recovered no ground-truth structure at any k (ARI vs. truth = 0) and cluster quality collapsed with granularity (silhouette 0.21 to 0.05), even though scDHA produced a moderately reproducible technical split. Two conclusions follow. First, the test is sensitive: it recovers real structure down to a 5% subpopulation and to weak (de = 0.03) separation. Second, and directly relevant to the rod result, a genuinely rare 5% subpopulation is recovered perfectly (ARI vs. truth = 1.00; cross-seed ARI = 1.00), so the rejection of the putative rare rod sub-state (cross-seed ARI 0.69) reflects a true absence of reproducible structure rather than a limit of detection.

**Figure 7.**
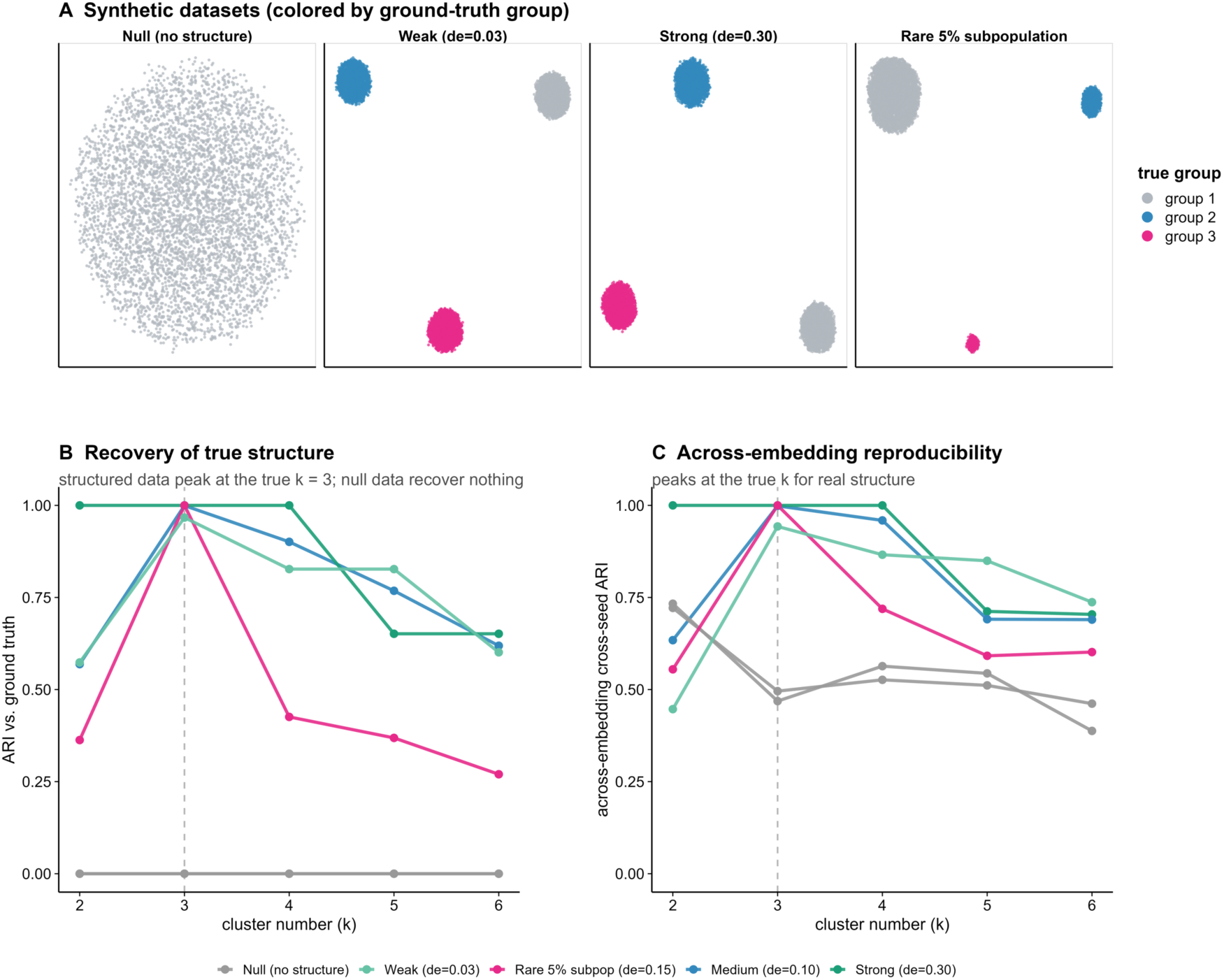
Ground-truth validation on synthetic data (Splatter). Null datasets contain one homogeneous population; structured datasets contain three groups at differential-expression probability de = 0.03 (weak), 0.10 (medium), and 0.30 (strong), plus a rare case with group sizes 80/15/5% (de = 0.15). Five independent scDHA embeddings per dataset. (A) UMAPs of representative simulated datasets colored by the true group: the null is one diffuse population, the structured datasets form distinct groups, and the rare dataset contains a small 5% subpopulation (pink). (B) ARI against the true labels versus cluster number k; all structured datasets peak at the true k = 3 (dashed line), including the rare 5% subpopulation and the weak-separation case, while null datasets recover no true structure at any k (ARI = 0). (C) Across-embedding reproducibility (cross-seed ARI) versus k, which peaks at the true k for real structure. The rare-subpopulation result confirms that a genuinely rare (5%) group is recovered perfectly, so the rejection of the putative rare rod sub-state reflects true absence rather than a detection limit.

### Pathway signatures of the metric-selected clusters

For completeness we characterized the selected k = 3 partition functionally. Cluster-defining differentially expressed genes (DEGs; Wilcoxon rank-sum, adjusted p < 0.05, |log₂ fold-change| > 0.25) numbered 1,460, 1,471, and 1,528 for the three clusters, respectively; all three clusters are recovered at every sampled timepoint (P15, P49, P75, P91). Submitting these gene lists to QIAGEN Ingenuity Pathway Analysis (IPA) (Kramer et al. 2014) returned distinct canonical-pathway profiles, chromatin/transcription terms for the dominant cluster and translation/ribosomal terms for the smaller clusters. Because this partition is not reproducible across embeddings, and because a high ribosomal/translation signature in a small cluster is a recognized technical signature in scRNA-seq, we interpret these enrichments as descriptive of one embedding’s partition rather than as evidence of defined biological rod sub-states; establishing a genuine rod sub-state would require an embedding-independent signal and orthogonal validation.

## Discussion

We present a label-free, multi-metric pipeline that reframes scRNA-seq clustering as an auditable decision-support problem: rather than nominating one algorithm as best, it reports five non-redundant quality metrics separately and asks whether subtype structure exists, at what granularity, and how reproducibly. Two design choices distinguish it from existing benchmarks (Duo et al. 2018; Freytag et al. 2018; Krzak et al. 2019; Germain et al. 2020): the metrics are reported individually so the analyst can see what each eliminates, and the framework is built to operate where ground truth is unavailable, with Harmony batch correction applied at both the whole-dataset and within-cell-type levels. The Phase 1 positive control is the load-bearing demonstration: the pipeline reproduces a hand-built eight-type annotation at ARI = 0.91 without labels, replacing the manual over-cluster-then-collapse workflow with an automated, deterministic benchmark.

The two discovery applications together validate the pipeline’s central claim on both a positive and a negative case. On bipolar cells, which have genuine subtypes, across-embedding reproducibility rose with cluster number to a plateau at the true subtype count (mean pairwise ARI to ∼0.93 at k ≈ 15, matching the ∼15 known bipolar types) with the quality metrics improving in parallel, the pipeline accepts real sub- structure at the correct granularity. On rods, the finer partition that the metrics select on any one embedding is not reproducible when the embedding is retrained (mean cross-seed ARI = 0.69; smallest cluster spanning 134–5,149 cells), and the metrics degrade with cluster number, the pipeline rejects over-clustering. Because both partitions pass a within-embedding bootstrap-stability check, the two cases would be indistinguishable by the resampling stability that is standard in the field; only the across- embedding test separates them. This is a cautionary result: a high bootstrap Jaccard, reported in isolation, would routinely be cited as proof of a reproducible rare sub-population. The practical lesson is that resampling stability and embedding-perturbation stability measure different things, and a claimed sub- population should be required to survive both. By surfacing this distinction automatically, the pipeline guards against a common and consequential form of over-interpretation while still recovering real sub- structure when it is present.

The rod result warrants a precise statement of what is and is not concluded. Because the pipeline recovers real subpopulations down to 5% of cells and to weak separation (de = 0.03) on synthetic data, the absence of a reproducible finer rod partition is an affirmative result, no discrete rod subpopulation of comparable abundance or separation is present, rather than a failure of sensitivity. It does not, however, exclude continuous or graded transcriptional variation among rods, which the high silhouette of the dominant- versus-minority split indicates is present; nor does it exclude structure that is too subtle to reach the detection threshold, or a state that is both extremely rare (well below 5%) and extremely weakly separated. The conclusion is specifically that rods do not contain discrete, reproducible subpopulations of detectable abundance and separation, and that a previously reported three-state rod decomposition reflected a single embedding rather than the data.

Several limitations apply. The five-metric panel informs but does not replace analyst judgment; its role is to expose the evidence, including disagreements among metrics. scDHA is non-deterministic across runs, the phenomenon the seed-robustness analysis quantifies; we cache embeddings for within-run reproducibility and report multi-seed sensitivity rather than a single run. The demonstration dataset has biological replicates only at P15 and P75, so per-timepoint abundance differences at single-replicate timepoints (P49, P91) cannot be cleanly separated from per-sample variation. Finally, although our synthetic validation (Fig. 7) recovers structured populations down to a 5% subpopulation and weak separation while rejecting null data, a more exhaustive sweep across the full continuum of subtype-size asymmetries and separation strengths would further delineate the method’s sensitivity limits.

In summary, a multi-metric, label-free, embedding-aware benchmarking pipeline recovers known retinal cell types without supervision and, applied to rod photoreceptors, identifies a single dominant transcriptional state while flagging that finer sub-structure is not reproducible across embeddings. The framework is applicable to any cell type or whole-dataset analysis as a reproducible alternative to single-method, single-embedding clustering, and its complete source code is publicly available (see Data access and Software availability).

## Methods

### Data acquisition and preprocessing

Six publicly available mouse-retina scRNA-seq datasets were obtained from the Gene Expression Omnibus (GEO): two biological replicates from postnatal day 15 (P15) (GSM8133387, GSM8133389; Yu Chen Lab), one P49 sample (GSM7734028; University of Oxford), two P75 samples (GSM7720305, GSM7720306; Johns Hopkins University), and one P91 sample (GSM6205478; Peking University First Hospital). All datasets were generated on the 10x Genomics Chromium platform and obtained as Cell Ranger output (matrix.mtx, features.tsv, barcodes.tsv), then converted to Seurat v5 objects (Hao et al. 2024). Doublets were detected with scDblFinder (Germain et al. 2021) and discarded. Cells were retained if they had 400–80,000 unique molecular identifier (UMI) counts, 400–8,000 detected genes, <18% reads mapping to mitochondrial genes, and <0.3% reads mapping to hemoglobin genes; mitochondrial and hemoglobin genes, and genes detected in fewer than five cells, were removed from the feature set.

Expression was normalized with SCTransform, regressing out the percentage of mitochondrial reads (Hafemeister and Satija 2019; Choudhary and Satija 2022). We computed 100 principal components and integrated samples with Harmony (Korsunsky et al. 2019) via Seurat’s IntegrateLayers function (theta = 2, lambda = 1, maximum iterations = 10). Louvain clustering was run at resolutions 0.2, 0.4, 0.6, 0.8, and 1.0; resolution 0.6 (34 clusters) was used for annotation. Cell types were assigned by cross-referencing cluster-level differentially expressed genes against canonical retinal markers. Rod photoreceptors were identified by *Rho* and *Nrl* expression.

### Rod isolation and embeddings

Rods were extracted from the integrated dataset and passed through a cone-contamination filter (module scores over canonical cone and rod marker panels; 61 cells removed), yielding 36,771 rods. For the linear embedding, raw counts were log-normalized, the 3,000 most variable genes were selected, the data were scaled, and PCA was run with 50 components. For the nonlinear embedding, scDHA (Tran et al. 2021) was applied to log(x + 1)-transformed counts (n = 3,000 genes; gene filtering enabled; seed = 1).

Harmony was then re-applied to each rod-level embedding using sample of origin as the batch covariate (theta = 2, lambda = 1, maximum iterations = 10). Both embeddings were cached and reused throughout the pipeline for within-run reproducibility.

### Clustering configurations and quality metrics

Seven configurations were evaluated: PCA + K-means, PCA + GMM (mclust), scDHA + K-means, PCA + Louvain, scDHA + Louvain, PCA + Leiden, and scDHA + Leiden. For graph-based methods, the resolution parameter was located by bisection followed by a logarithmic-grid fallback to reach each target k; the achieved k is reported alongside the target k, and cells for which the sweep could not reach the target k are masked in the heatmaps. Each (method, k) was scored on five non-redundant metrics: (i) ROGUE (Liu et al. 2020), within-cluster transcriptomic homogeneity anchored to the unsplit-population baseline; (ii) cluster-size balance = 1 − G, where G is the Gini coefficient of cluster sizes; (iii) per-cluster bootstrap Jaccard (Jaccard 1912; Hennig 2007), the minimum across clusters of the mean maximum Jaccard between each original cluster and any bootstrap cluster over resamples; (iv) marker specificity, the mean over clusters of (pct.1 − pct.2) × |log₂ fold-change| for significant positive markers from FindAllMarkers; and (v) silhouette width (Rousseeuw 1987). For the Phase 1 positive control we additionally computed the ARI (Hubert and Arabie 1985) and NMI against the manual annotation; these are post-hoc and are not used to select a partition.

Two |log₂ fold-change| thresholds are used and serve different purposes. The marker-specificity metric uses the stringent |log₂ fold-change| > 0.5 (only positive markers, minimum detection fraction 0.25) appropriate for a specificity score, whereas the cluster-defining DEGs reported for the chosen partition use the more permissive |log₂ fold-change| > 0.25 standard for DEG discovery.

### Stability analysis

Two complementary stability analyses were performed on the chosen partition. Within-embedding stability was quantified by 50 bootstrap re-clusterings (80% subsampling) of the fixed Harmony-corrected scDHA embedding, reporting the per-cluster Jaccard and the bootstrap-to-reference ARI. Across- embedding stability was quantified by re-training scDHA with five independent random seeds on the identical input, re-applying Harmony, re-clustering, and computing the pairwise ARI/NMI between the resulting partitions. For the discovery-mode controls (bipolar cells and rods), this across-embedding reproducibility was computed as a function of cluster number (bipolar cells to k = 18 to span the ∼15 known subtypes; rods to k = 9) and reported alongside the silhouette, ROGUE, and balance metrics at each k, so that the concordance between reproducibility and cluster quality could be assessed. Adjusted Rand index and NMI were computed with the aricode package (Chiquet et al. 2020).

### Synthetic validation

Synthetic scRNA-seq datasets (5,000 cells, 2,000 genes) were generated with Splatter (Zappia et al. 2017). Null datasets were simulated as a single population; structured datasets were simulated with three groups (splatSimulateGroups) at differential-expression probabilities de = 0.03, 0.10, and 0.30, and a rare- subpopulation dataset with group proportions 0.80/0.15/0.05 (de = 0.15). Each dataset was processed through five independent scDHA embeddings, and at each k = 2–6 we recorded across-embedding reproducibility, the ARI against the known simulated labels, and the silhouette width, using the same procedures as for the real cell types.

### Pathway analysis

Cluster-defining DEGs from the chosen partition (Wilcoxon rank-sum, adjusted p < 0.05, |log₂ fold-change| > 0.25, positive only) were submitted to QIAGEN Ingenuity Pathway Analysis (IPA) (Kramer et al. 2014). Canonical pathways and predicted activation z-scores were retrieved. Because IPA significance scales with input gene-list size, absolute −log₁₀(p) magnitudes are not compared across clusters of different size; the direction of the z-scores is the primary readout.

### Statistics, software, and reproducibility

Analyses used R 4.5.3 with Seurat v5 (Hao et al. 2024), scDblFinder (Germain et al. 2021), scDHA (Tran et al. 2021), ROGUE (Liu et al. 2020), mclust (Scrucca et al. 2016), harmony (Korsunsky et al. 2019), igraph, FNN, and aricode (Chiquet et al. 2020). Random seeds were fixed wherever supported; scDHA is non-deterministic across runs, and this residual variability is itself quantified by the across-embedding stability analysis. The complete pipeline is implemented as modular R scripts and runs end-to-end from the raw 10x Genomics inputs.

### Software availability

The complete, deterministic pipeline (stages 01–06 plus the two stability stages) is available at https://github.com/zachyousef/senior-design-pipeline (made public upon acceptance) and as Supplemental Material.

### Data access

All scRNA-seq datasets analyzed here are publicly available from the Gene Expression Omnibus under accession numbers GSM8133387, GSM8133389, GSM7734028, GSM7720305, GSM7720306, and GSM6205478. No new sequencing data was generated, and no animal or human procedures were performed for this study; all data are secondary analyses of previously published, publicly deposited datasets. Processed outputs (cluster assignments, metric tables, and stability results) are provided in the repository below.

### Use of AI tools

Claude (Anthropic) was used to assist with code organization and debugging. All content was reviewed, verified, and edited by the authors, who take full responsibility for the accuracy and integrity of this work.

## Acknowledgments

The authors acknowledge the Office of Advanced Research Computing (OARC) at Rutgers, The State University of New Jersey for providing access to the Amarel cluster and associated research computing resources that have contributed to the study reported here. URL: https://it.rutgers.edu/research-computing

## Author contributions

Z.Y. designed and implemented the pipeline, performed the analyses, and wrote the manuscript. J.S., D.K., A.B., and P.B. contributed to data curation, pipeline development, and analysis. H.C. contributed to design and manuscript preparation. J.W. and H.W. contributed to data analysis, discussion, and manuscript revision. L.C. supervised the study. All authors reviewed and approved the manuscript.

